# A population MRI brain template and analysis tools for the macaque

**DOI:** 10.1101/105874

**Authors:** Jakob Seidlitz, Caleb Sponheim, Daniel Glen, Frank Q. Ye, Kadharbatcha S. Saleem, David A. Leopold, Leslie Ungerleider, Adam Messinger

## Abstract

The use of standard anatomical templates is common in human neuroimaging, as it facilitates data analysis and comparison across subjects and studies. For non-human primates, previous *in vivo* templates have lacked sufficient contrast to reliably validate known anatomical brain regions and have not provided tools for automated single-subject processing. Here we present the “*National Institute of Mental Health Macaque Template”*, or NMT for short. The NMT is a high-resolution *in vivo* MRI template of the average macaque brain generated from 31 subjects, as well as a neuroimaging tool for improved data analysis and visualization. From the NMT volume, we generated maps of tissue segmentation and cortical thickness. Surface reconstructions and transformations to previously published digital brain atlases are also provided. We further provide an analysis pipeline using the NMT that automates and standardizes the time-consuming processes of brain extraction, tissue segmentation, and morphometric feature estimation for anatomical scans of individual subjects. The NMT and associated tools thus provide a common platform for precise single-subject data analysis and for characterizations of neuroimaging results across subjects and studies.

## Introduction

Investigations into the structure and function of the non-human primate brain significantly contribute to our overall understanding of the nervous system. The macaque monkey is a well-studied model system that has provided tangible translational benefits, owing to its phylogenetic proximity to humans (Zhang et al., 1993) and the ability to test hypotheses using invasive techniques (e.g., electrophysiology, histology, and lesions). The application of non-invasive brain imaging techniques, such as structural and functional magnetic resonance imaging (MRI), in both humans and monkeys has helped contextualize findings from human research and demonstrate the translational relevance of the macaque as a model system. However, to reap the most translational benefit from non-human primate neuroimaging, it is essential that the analytic tools used in monkey imaging keep parity with the tools used in human imaging and that these tools be made widely available.

In MRI research, multi-subject analysis bolsters scientific validity by increasing statistical power and highlighting reliable neurological phenomena across a population (Friston et al., 1999). To facilitate comparison across subjects, data from each subject is typically transformed to a common image of the brain’s anatomy, with an associated coordinate space, for visualization and analysis (Holmes et al., 1998; Talairach & Tournoux, 1988). This anatomical template is often an individual subject’s brain, such as the Colin N27 brain (Holmes et al., 1998). Others, such as the Montreal Neurological Institute’s ICBM152 template (Mazziotta et al., 2001), are averages of the anatomies of multiple individuals. In principle, such multi-subject templates are preferable for group-level analysis because they possess features that are typical of the population’s brain anatomy and thus have greater cross-subject validity. In practice, the quality of multi-subject templates depends on how well the individual brains are registered (*i.e.*, aligned) prior to averaging.

Templates are less commonly used in monkey MRI research for two reasons. First, macaque neuroimaging studies typically involve a small number of animals, so multi-subject analysis is limited. Second, existing T1-weighted templates have either been based on a single animal or lacked sufficient detail for precise anatomical localization. Single-subject templates reflect the idiosyncratic anatomy of an individual, rather than the species as a whole (Reveley et al., 2016; Van Essen et al., 2001). Multi-subject templates may better reflect macaque brain anatomy. However, templates based on linear registration methods (Black et al., 2004; McLaren et al., 2009) have produced blurry averages, making anatomical localization difficult. Templates based on nonlinear transformation techniques have displayed improved detail and contrast, but not to the extent of a recent single-subject *ex vivo* template (Reveley et al., 2016).

We sought to create an improved and representative *in vivo* macaque template. To do so, we scanned a large cohort of animals at high field-strength, and then nonlinearly and iteratively averaged these scans using a validated template-creation process (Avants et al., 2010). This process does not favor any one individual, but rather represents an unbiased average of the population used to create it (Avants et al., 2010). The resulting template, which we call the National Institute of Mental Health Macaque Template - or NMT for short - contains emergent anatomical details not evident in either the individual scans used to create it or in previous *in vivo* templates. To take full advantage of the NMT’s representative nature, we have segmented its different tissue types (and created corresponding surface reconstructions) and generated a map of its cortical thickness. We are making the NMT volume, tissue segmentation, and surface representations openly available to the research community. In addition, we are providing accompanying tools for automated single-subject analysis.

The NMT package will give a broad range of researchers (within and outside of neuroimaging) a high-resolution platform and standardized coordinate system for localization and visualization of any spatially distributed brain-related data. The tools we provide will streamline analysis of both single- and multi-subject MRI data, which will allow for robust cross-animal comparison and foster collaboration across research groups and institutions.

## Materials and Methods

### 2.1 Subject Information

Our subject cohort consisted of 31 rhesus macaques (*Macaca mulatta*) from the Central Animal Facility at the NIMH. The monkeys were juveniles and adults between 3.2 and 13.2 years old when the anatomical scans were collected (average of 5.5 years). The ages of the 25 males and 6 females were comparable (5.6 ± 2.3 and 5.3 ± 1.1 years, respectively). The monkeys weighed 6.5 kg on average at the time of scan collection, with the males weighing significantly more than the females on average (6.9 ± 1.9 vs. 5.0 ± 0.8, one-tailed t-test, *p* < 0.0004). All animals were under food restriction on the date of scan collection. Prior to the date of the scan, none of the monkeys scanned had ever undergone an invasive brain procedure (e.g., craniotomy). All animal procedures were conducted in compliance with the National Institutes of Health Guide for the Care and Use of Laboratory Animals.

### 2.2 Scanning Protocol

T1-weighted MR anatomical images were acquired in a 4.7 T horizontal scanner (Bruker Biospec 47/40) using a modified driven equilibrium Fourier transform (MDEFT) method (Lee et al., 1995; Deichmann et al., 2004) at the Magnetic Resonance Imaging Facility at the NIH. Each macaque was anesthetized with isoflurane and placed into the scanner in a sphinx position with its head secured in a holding frame. A single loop circular coil, with a diameter of either 14 or 16.5 cm, was placed on top of the animal’s head. The whole-brain MDEFT images were acquired in a 3D volume with a field of view of 96 × 96 × 70 mm^3^, and 0.5 mm isotropic voxel size. Each MDEFT scan was acquired over 40-60 minutes. Typically, several MDEFT scans were collected consecutively (average = 2.5 scans/monkey, range 1-7 scans). These scans were then rigidly aligned to the first and averaged within subject. This within-subject average was used as the input for the template creation process, so that one anatomical image was included for each subject (Figure 1, Step 0).

**Figure 1.**
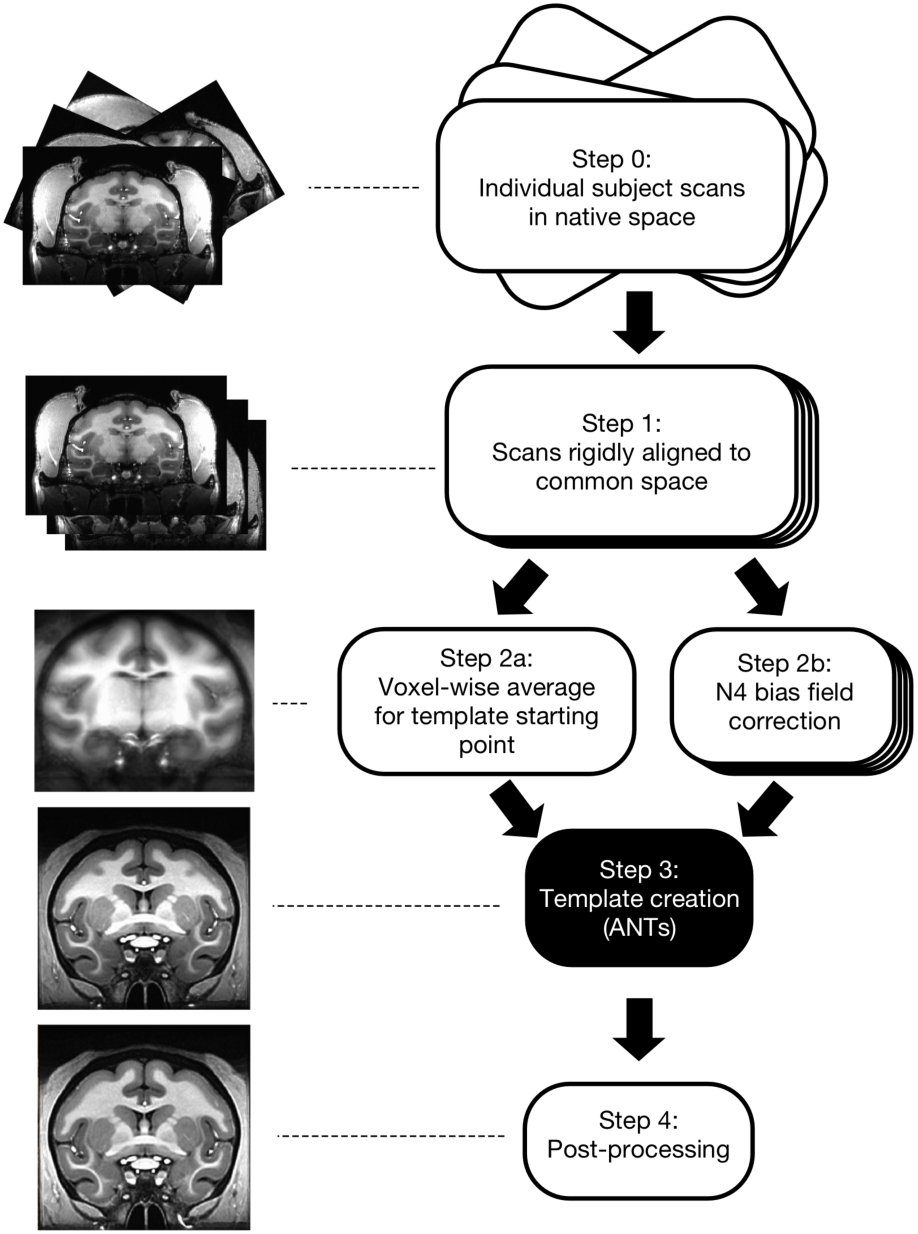
Pipeline for creating the NIMH Macaque Template (NMT). Step 0) T1-weighted MRI scans of the brain were collected from 31 macaque monkeys *in vivo* and averaged within subject. Step 1) These anatomical scans were aligned to that of an independent macaque monkey using a 6-parameter rigid-body transformation. Step 2a) A voxel-wise average was calculated across the aligned normalized subject images, forming the initial reference template for Step 3. Step 2b) Each subject’s aligned volume was corrected for non-uniformities in intensity values. Step 3) Each subject’s volume was affine-transformed to the current reference template using 12 parameters. Voxel-wise nonlinear (diffeomorphic) transformations were then calculated from each subject’s anatomy to the current reference template. For successive iterations, reference templates were updated by averaging the aligned subject volumes from the previous iteration. Then, a scaled average of the inverse affine and diffeomorphic transformations across subjects was applied to this updated template. This procedure was iterated four times. Step 4) The output template from Step 3 was rigidly aligned to be in the orientation shown in Figure 2 and capped at a maximum intensity value. A mask of the brain was generated, and the volume within this brain mask was corrected for non-uniformities in intensity values.

### 2.3 Template Creation

Image analysis was performed using the freely available software packages AFNI (<u>https://afni.nimh.nih.gov/afni/</u>; Cox, 1996) and ANTs (<u>picsl.upenn.edu/software/ants/</u>; Avants et al., 2010). To increase data processing efficiency, we made use of the cluster-computing and parallelization capabilities available through the computational resources of the NIH HPC Biowulf cluster (hpc.nih.gov). The main steps for the template creation process are shown in Figure 1. Whole-head images were used so that the template would accurately represent the brain-skull boundary (Step 0; Scott et al., 2015). Initially, subjects were coarsely aligned and resampled from 0.5 to 0.25 mm isotropic resolution (Step 1). A 6-parameter rigid-body transformation was used to align each of the 31 subject images to an independent coordinate space (subject D99-S from Reveley et al., 2016). The voxel-wise average of these subject images served as the initial target image in the template creation process (Step 2a). Then, we applied N4 bias field correction (Avants et al., 2011) separately to each of the aligned subject images to normalize variations in image intensity across each volume (Step 2b). To create the population-average template, we used symmetric group-wise normalization (SyGN), an iterative nonlinear registration process (Avants et al., 2010; Love et al., 2016). This was carried out using the ANTs *buildtemplateparallel.sh* script (Step 3), described as follows: each subject’s brain was aligned to the current target image via a 12-parameter affine transformation and a nonlinear (diffeomorphic - allowing for local warps in structure) transformation. These aligned images were averaged to generate an improved template image. The inverse of the affine and diffeomorphic transformations was averaged across subjects, scaled, and applied to this template image to align it closer to the original input anatomies. This process was iterated, with the updated template image serving as the new target image for registration with the original subject images, until convergence between successive target images occurred. The output image from the script lies intermediate to all the input anatomies and, thus, is not biased towards any single individual. See section 2.5 for a detailed description of the post-processing (Step 4) performed on the output template.

### 2.4 Contrast-to-noise Ratio

The contrast-to-noise ratio (CNR) measures how distinguishable different tissue classes are from one another. Here, CNR is defined as the mean intensity of white matter (WM) minus the mean intensity of the gray matter (GM) divided by the standard deviation of the intensities in the cerebrospinal fluid (CSF). These values were calculated over spherical ROIs (radius = 0.5 mm, N = 33 voxels) located in the corpus callosum for WM, left and right caudate nuclei for GM, and superior sagittal sinus for CSF. These ROIs were all centered on a coronal slice at the rostral-most point of the anterior commissure (AP +1). For each subject, CNR was calculated on the subject’s N4-corrected image nonlinearly aligned to the template using these same ROIs. The CNR calculation for the template was computed before the post-processing steps described below.

### 2.5 Post-processing, Segmentation, and Surface Generation

The intensity values in the volume produced by the template creation script were capped to account for the extreme values of the blood vessels, thus preventing the need for manual adjustment of standard image viewers. This qualitatively improved the subsequent N4 bias field correction (Tustison et al., 2010) and the accuracy of the intensity-based brain extraction. Brain mask creation was performed using the MIPAV software suite (McAuliffe et al., 2001), which supplies a modified version of FSL’s Brain Extraction Tool (Smith, 2002). Although the outputted brain mask was highly accurate, manual editing was performed using MIPAV to correct the mislabeling of any non-brain matter as brain, and any brain matter as non-brain. N4 bias field correction was performed over just the volume within the edited brain mask to correct for any remaining non-uniformities in image intensity.

This post-processed NMT volume was segmented by *k*-means clustering (*k* = 3 classes) using the *Atropos* command in ANTs (Avants et al., 2011). This produced separate probabilistic tissue segmentation maps for the three classes, which from brightest to darkest roughly corresponded to WM, GM, and CSF. Surface reconstructions of WM and GM were generated from their respective segmentation maps (thresholded at 50% probability and manually edited), using AFNI’s *IsoSurface* and *Surfsmooth* commands (Lewiner et al., 2003), and inflated using the freely available Connectome Workbench software (Figure 5; Marcus et al., 2011).

### 2.6 NMT Coordinate Space

The NMT was manually translated and rotated to match the Horsley-Clarke stereotaxic plane (Schurr & Merrington, 1978), which is the standard orientation used during stereotaxic surgeries. This Horsley-Clarke stereotaxic plane contains the intra-aural meatus and infraorbital ridge in the same horizontal slice. Because these structures were not within the field of view of the template (or many of the individual subjects), we approximated this orientation by aligning the superior aspect of the anterior commissure with the inferior aspect of the posterior commissure (Saleem & Logothetis, 2012). To maintain a similar coordinate space as the D99-S template (Reveley et al., 2016), we set the origin of NMT coordinate space to be the center of the anterior commissure (see Figure 2). The NMT is defined on a 253 × 339 × 241 voxel grid, with a 0.25 mm isotropic voxel size. Like the D99-S template, the NMT grid is organized with RAI (“Neurological”) index ordering (i.e., × = medial-to-lateral, y = anterior-to-posterior (AP), z = superior-to-inferior (SI); see Figure 2).

**Figure 2.**
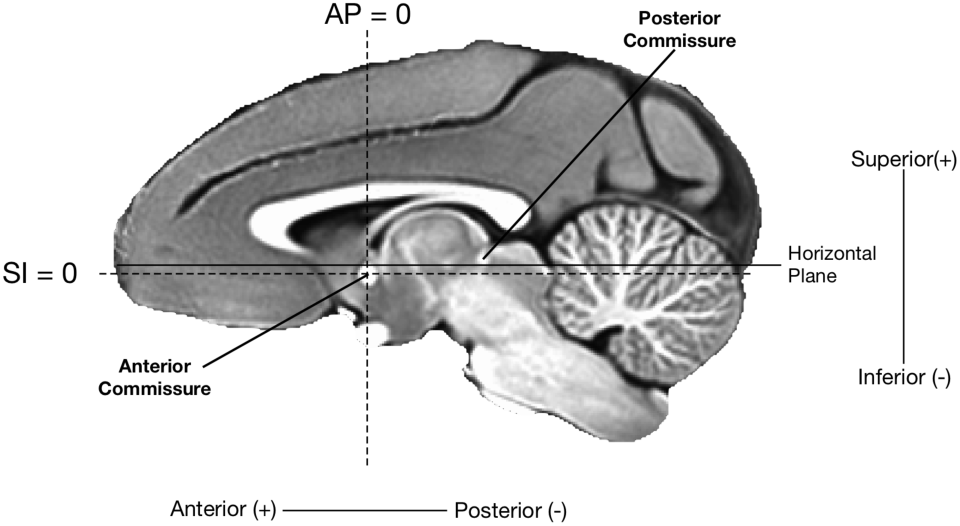
The orientation and coordinate system of the template. The NMT was rotated so that the dorsal-most point of the anterior commissure and the ventral-most point of the posterior commissure (solid line) lay in the horizontal plane. This approximately aligns the NMT with the Horsley-Clark stereotaxic coordinate system, whose horizontal is defined by the infraorbital ridge and the line connecting the centers of the external auditory meati (Horsley & Clark, 1908; Saleem & Logothetis, 2012). In keeping with previous anatomical templates, the origin of the NMT was defined to be at the center of the anterior commissure (AP 0, SI 0, intersection of the dashed lines), midway between the hemispheres (ML 0). The sagittal slice shown is one millimeter to the right of the midline (ML +1).

### 2.7 Morphological Variability across the NMT Subject Cohort

We assessed the variability in morphology across the cohort by calculating the voxel-wise Mean Positional Difference (MPD; Calabrese et al., 2015; Kovacevic et al., 2005). For each subject, MPD was calculated as the average of the absolute values of the diffeomorphic warp vector fields (i.e., spatial displacements along each of the three axes) at each voxel from the 3-dimensional diffeomorphic warp files outputted during the template creation process. These values were then averaged across subjects to generate the whole-brain MPD map. Smaller MPD values meant that the morphology of many or all of the subjects was in close agreement with the NMT average.

### 2.8 NMT Cortical Thickness

We estimated the cortical thickness (CT) of the NMT, in part, to validate the NMT against previous templates that have reported this metric. Classic surface-based CT algorithms, such as that used in FreeSurfer (Fischl & Dale, 2000), require the GM surface (boundary between GM and CSF) and WM surface (boundary between GM and WM) to have the same number of vertices, so that the Euclidean distance can be calculated between each set of corresponding vertices. However, the GM and WM surfaces of the NMT were generated independently to increase their precision. Consequently, because the surface vertices were unmatched, we used a volumetric approach to estimating CT. To create a GM volume for this process, we thresholded the GM probability map (see section 2.4) at 50% and manually edited this mask to correct clear misclassifications of tissue (e.g., WM in the extreme capsule and occipital cortex that was erroneously classified as GM). CT was estimated over this edited GM volume using the *KellyKapowski* command in ANTs, which is based on the DiReCT method (Das et al., 2009). This produces a thickness estimate for each voxel in the GM volume. CT generated by this volume-based method and by surface-based algorithms have been shown to yield similar results (Tustison et al., 2014).

## Results

### 3.1 The NIMH Macaque Template

Figure 3a shows select axial slices of the NIMH Macaque Template (NMT). Importantly, the template includes not only the brain (shown in red) but also the other parts of the head, including skull, muscles, and eyes, for reference. The NMT has high contrast (CNR = 12.71) and clear edges, making it easy to differentiate the brain from other structures and to distinguish the borders between both cortical and subcortical gray matter and white matter. The contrast of the NMT was greater than that of all but one of the MRI scans of the individual monkeys used to create it (mean CNR = 6.75 ± 0.52, N=31, std. err). The transitions between tissue types in the NMT were sharper than in individual monkey scans (see Figure S1) and previous *in vivo* multi-subject templates (see section 3.6 and Figure 4). The NMT’s clarity allows for assessment of detailed structural brain morphology and for anatomical localization of both neuroimaging and non-imaging data.

**Figure 3.**
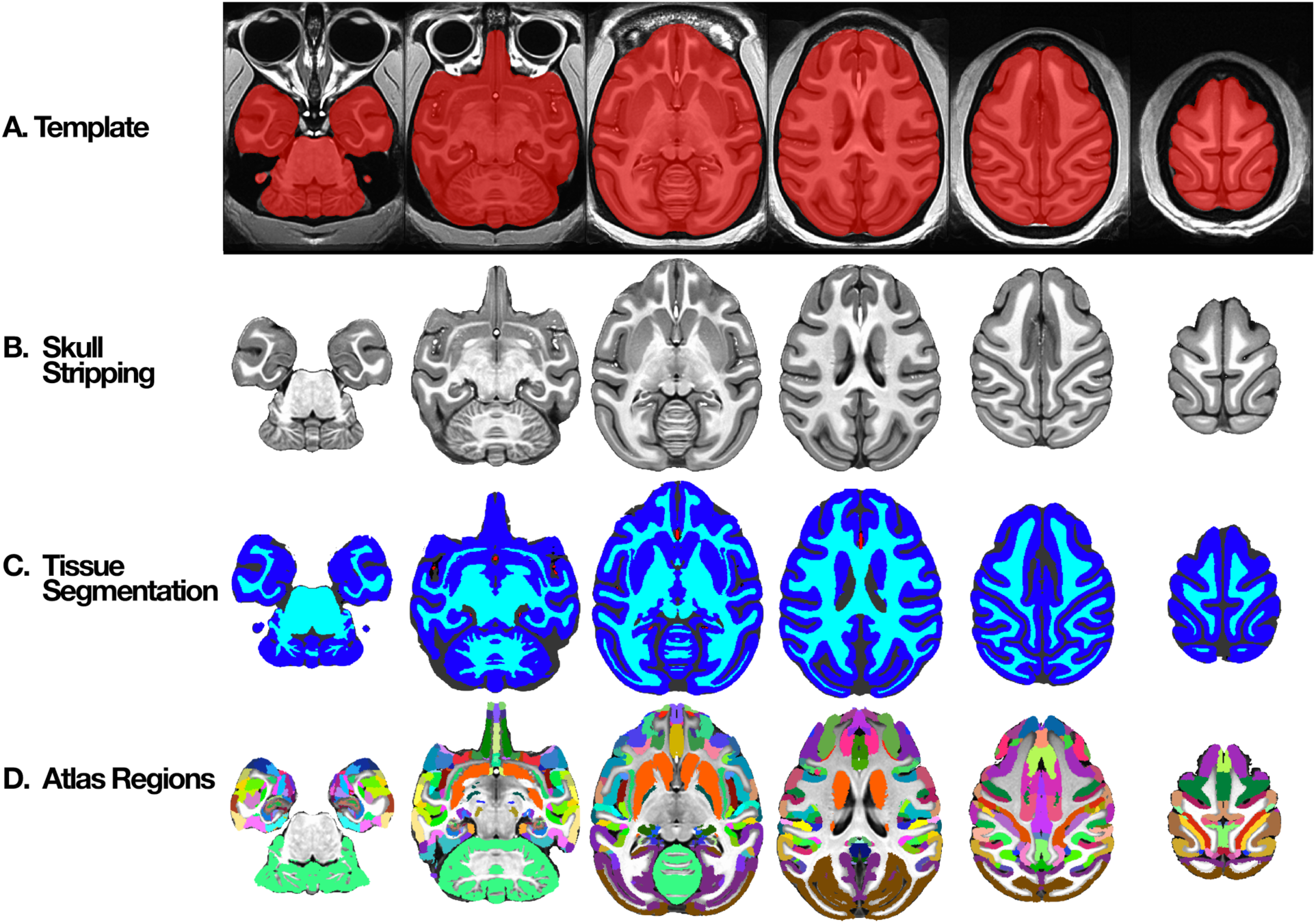
Axial slices through NMT with segmentation. A) Whole head template with brain mask (red). B) Extracted brain template. C) Segmented, manually-corrected tissue classes – gray matter (GM; dark blue), white matter (WM; cyan), cerebrospinal fluid (CSF; dark gray), and blood vasculature (BV; red). In addition to the tissue masks, the NMT distribution also includes probabilistic tissue segmentation maps of the GM, WM, and CSF. D) An example of nonlinear alignment of a digital anatomical atlas (Reveley et al., 2016; Saleem & Logothetis, 2012) to the NMT.

### 3.2 Brain Mask

The clear borders of the NMT greatly simplified the generation of a brain mask for distinguishing the brain from surrounding tissue. Automated processes for identifying brain tissue are unreliable at best when applied to monkey MRI scans. As a result, the process of segmenting the brain is typically labor intensive. In the case of the NMT, however, an automated process (MIPAV) was largely successful. We made small manual corrections to the mis-identified and unidentified brain tissue in this initial mask to produce the brain mask shown in red in Figure 3a. The mask is largely complete, including the entire cerebellum and much of the brain stem. Parts of the central nervous system not included in the brain mask are the optic nerve, retina, and olfactory bulb. The brain mask includes portions of the dura mater as well as blood vessels within the brain. In addition to the brain mask, manually drawn masks of the olfactory bulb and cerebellum are provided separately.

Figure 3b shows the portion of the NMT within the brain mask (*i.e.,* after “skull stripping”). This image shows the clear differences in image intensities between the CSF, GM, and WM. This contrast is evident not only in the cerebral hemispheres but also in the cerebellum. Many subcortical structures can be differentiated as well (see section 3.6 below).

### 3.3 Tissue Classification

We classified the tissue within the brain mask into three categories, comprised predominantly of WM, GM, and CSF. The NMT’s high contrast allowed for automated tissue classification using *Atropos* in ANTs. The result is a probabilistic map showing the probability that each voxel belongs to each of the three tissue categories. We also generated separate binary masks of GM, WM, and CSF by assigning each voxel to its most probable tissue type (probabilistic maps thresholded at 50%). We manually edited these thresholded masks within parts of the cerebral cortex (e.g., claustrum, primary visual cortex [V1], frontal pole, ventral temporal lobe) to improve parcellation. Additionally, we manually defined the major arterial blood vasculature (BV) in the NMT volume (see Figure 7 and Figure 3c). This data represents the first population-level mapping of macaque vasculature. The BV is included as a fourth tissue category in the combined segmentation map (limited to within the brain mask). Separate surface and volumetric representations of the BV are included in the NMT distribution.

Figure 3c shows the manually edited tissue classification map. The GM mask delineates the cortex and some of the larger subcortical nuclei (e.g., striatum – the caudate and putamen). This mask includes small amounts of white matter directly adjacent to subcortical gray matter structures and small amounts of dura mater. The WM mask consists of white matter, as well as some subcortical structures (e.g., globus pallidus and other parts of the basal ganglia, parts of the thalamus and hypothalamus) that were not captured by the GM mask. The GM mask was further edited to remove all subcortical structures, resulting in a separate cortical GM mask. This cortical GM mask does not include two allocortical structures, namely hippocampus and the periamygdaloid cortex. The separate probabilistic tissue maps (GM, WM, and CSF) and single combined segmentation map (GM, WM, CSF, and BV) are included in the NMT distribution, enabling data analysis (e.g., clustering) within distinct tissue types.

### 3.4 Size of the NMT

The NMT brain (Figure 3b) is, at its largest extent, 73 mm in the anterior-posterior dimension, 58 mm in the medial-lateral dimension (both left and right hemispheres together), and 45.75 mm in the dorsal-ventral dimension. The total intracranial volume of the brain mask (Figure 3a) is 91.76 cc, of which the cerebellum is 7.27 cc. These measurements are consistent with brain volume analyses across large macaque populations (Franklin et al., 2000; Rilling & Insel, 1998; Scott et al., 2015). Using the 4-tissue segmentation mask, this total volume is composed of a GM volume of 50.29 cc, a WM volume of 28.81 cc, a CSF volume of 11.76 cc, a BV volume of 0.15 cc, and a volume of 0.74 cc for tissue not classified as one of the above. The GM volume is a slight underestimate (and the WM is a slight overestimate) because the WM mask includes some subcortical nuclei. The volume of the cortical GM is 38.5 cc. The olfactory bulb, located outside the brain mask, has a volume of 0.09 cc.

### 3.5 Regional Parcellation of the NMT

To facilitate seamless integration with previous work and analyses, we are supplying the nonlinear transformations to and from some published templates. For example, the transformations between the the single-subject F99 template and the NMT will allow users to take advantage of the regional parcellations and surface-based atlases derived for the F99 anatomy and available through the Caret software package (Van Essen et al., 2011). Similarly, transformation between NMT and the single-subject D99-S brain (Reveley et al., 2016) will allow users to avail themselves of the digital version of the Saleem & Logothetis (2012) atlas, which is available through the AFNI website. Figure 3d shows the anatomical parcellation of this digital atlas after nonlinear transformation to the NMT brain (Reveley et al., 2016). There is a high degree of alignment between the atlas areas and the underlying cortical and subcortical gray matter of the NMT brain.

### 3.6 Comparison of the NMT with Existing Macaque Templates

The diffeomorphic transformations used to create the NMT align anatomical features shared amongst individual animals. Thus, the NMT has sharp borders and high contrast between tissue classes, resulting in precise depiction of fine anatomical structures (Figure 4). The anatomical details are especially evident outside the neocortex. For example, in Figures 2, 3b, and 7b the individual folia of the cerebellum are visible. Figure 4A shows a coronal slice through the NMT, revealing several subcortical structures, including the claustrum, basal ganglia, and thalamus. The detail evident in this slice is an emergent property of the NMT that was not present in the individual scans (Figure S1) used in its generation.

**Figure 4.**
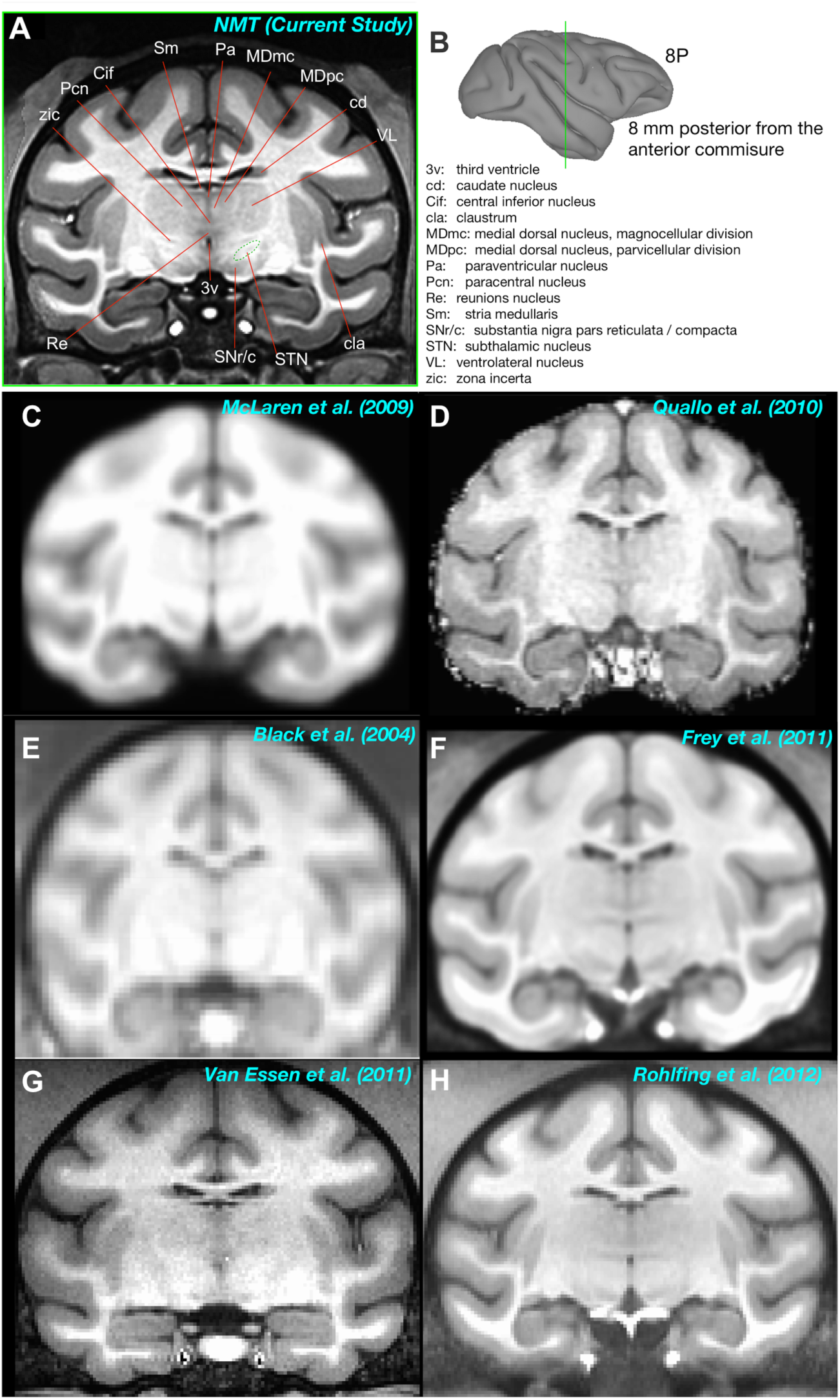
Comparison of the NMT to other available *in vivo* templates. A) Anatomical labeling of subcortical structures in an example coronal slice of the NMT. B) The location of the slice is depicted by the green line through the gray matter surface. The slice is 8 mm caudal to the anterior commissure (AP -8). Note that fine subregions of the thalamic nuclei and other subcortical structures (e.g., claustrum) are visible in the NMT that are not evident in previous single- and multi-subject *in vivo* templates (C-H).

Figure 4C-H shows approximately the same coronal slice through some previous *in vivo* templates of various macaque species. The templates that involved only affine transformations to align individuals before averaging (Figure 4C-E) are quite blurry because of differences between individual animals. This blurring remains whether the averaging is across 12 subjects (Figure 4E) or 112 subjects (Figure 4C). There is more detail in the F99 template, which is a scan from a single rhesus monkey (Figure 4G) and in the multi-subject templates that employed nonlinear registration methods (Figure 4F and H). But even in these cases, the border between the GM and WM is not as sharp as the NMT and only large subcortical structures, such as the thalamus and putamen, are evident.

In the NMT, on the other hand, one can delineate many thalamic subnuclei (e.g., ventrolateral nucleus, medial dorsal nucleus, and midline structures, such as reunions and paracentral nucleus), subthalamus (e.g. subthalamic nucleus and zona incerta), and the basal ganglia (e.g., caudate and putamen). One can also appreciate other small structures such as dura mater and arterial blood vessels. The fine detail present in the NMT will help accurately identify the brain areas responsible for a signal (e.g., an fMRI activation). This detail will also help constrain and improve the accuracy of the registration of individual subjects or cohorts (e.g. lesion groups) to the NMT.

### 3.7 NMT Surfaces

The manually edited tissue classification masks discussed above (section 3.3) were smoothed and converted into separate surfaces of the left and right hemispheres (as well as a distinct cerebellum surface, shown in Figure 7a, left). These surfaces are useful for concisely depicting data across the cortex and for use with surface-based tools, such as the surface-based macaque atlases in Caret (Van Essen et al., 2001). Figure 5 shows the “gray matter surface”, which sits at the border of the GM and CSF (also referred to as the “pial surface”). The surface representing the border between the GM and WM is labeled “white matter surface” in Figure 5. Inflated surfaces, often useful for visualization of sulci (Fischl et al., 1999), were created using Connectome Workbench (Marcus et al., 2011), and are also included in the data distribution. An approximated “mid-cortical surface” was generated by expanding the white matter surface with *mris_expand* (Fischl et al., 1999) by 1.0 mm, while controlling for intersections in boundaries. It is important to note that a surface representation only shows the data associated with the voxels that intersect that surface. As such, we suggest using the “mid-cortical surface” to depict volumetric data across the cortical surface.

**Figure 5.**
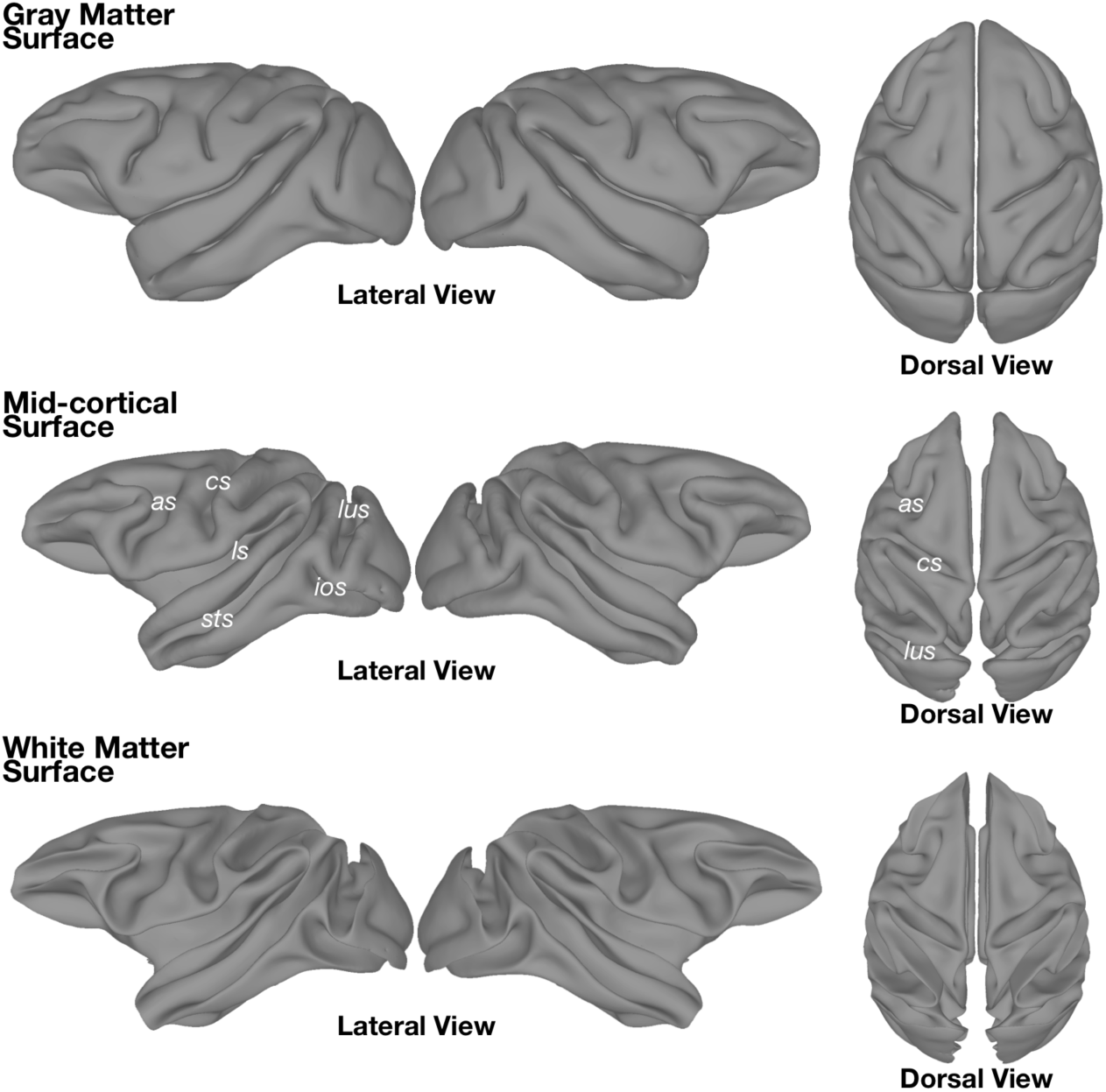
Cortical surfaces manually constructed from the segmentation of the NMT. Surface representations based on the boundary between (Top) gray matter and CSF, and (Bottom) gray matter and white matter. (Middle) The mid-cortical surface was generated by expanding the white matter surface 1 mm. The mid-cortical surface is recommended for surface visualization of volumetric data. All surfaces were artificially separated down the mid-sagittal plane to allow hemispheres to be viewed and inflated individually. The surfaces depicted, as well as their inflated versions, are included with the NMT distribution. *as* = arcuate sulcus; *cs* = central sulcus; *ios* = inferior occipital sulcus; *ls* = lateral sulcus; *lus* = lunate sulcus; *sts* =superior temporal sulcus.

**Figure 6.**
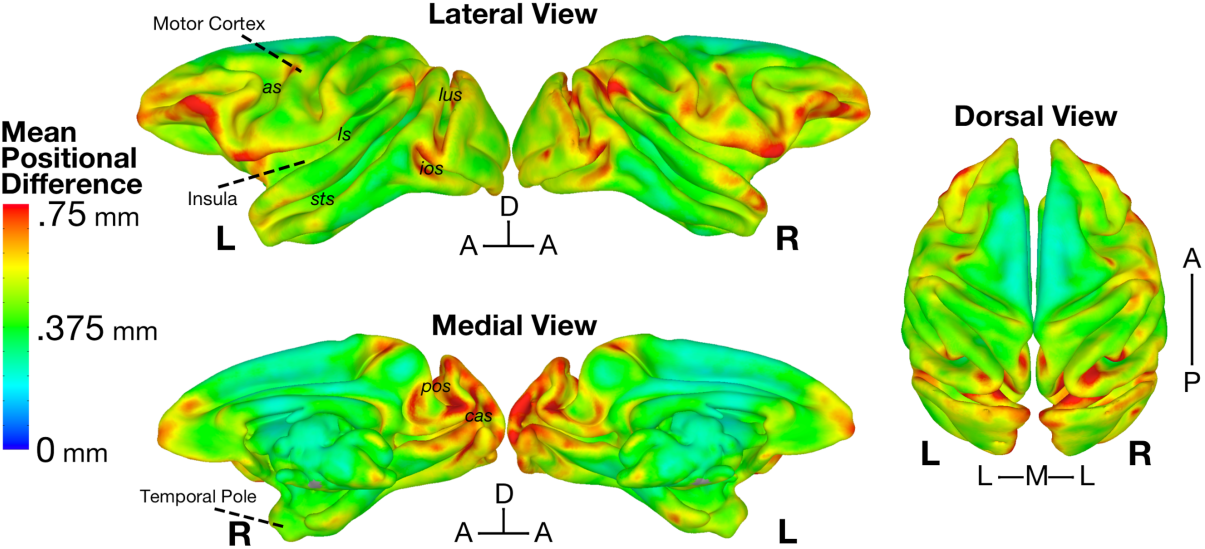
Average deformation between each subject and the NMT. The Mean Positional Difference (MPD) map shown on the mid-cortical surface of the NMT. The MPD at each voxel is the average absolute displacement of each subject’s diffeomorphic transformation. On average, voxels were nonlinearly warped by less than the dimension of a structural voxel (mean MPD = 0.45 mm). MPD was greatest in ventral prefrontal, frontal pole, ventral premotor, opercular, dorsal temporal pole, and parieto-occipital areas, which are known to be highly variable in their cortical folding. *as* = arcuate sulcus; *cas* = calcarine sulcus; *cs* = central sulcus; *ios* = inferior occipital sulcus; *ls* = lateral sulcus; *lus* = lunate sulcus; *pos =* parieto-occipital sulcus; *sts* = superior temporal sulcus.

**Figure 7.**
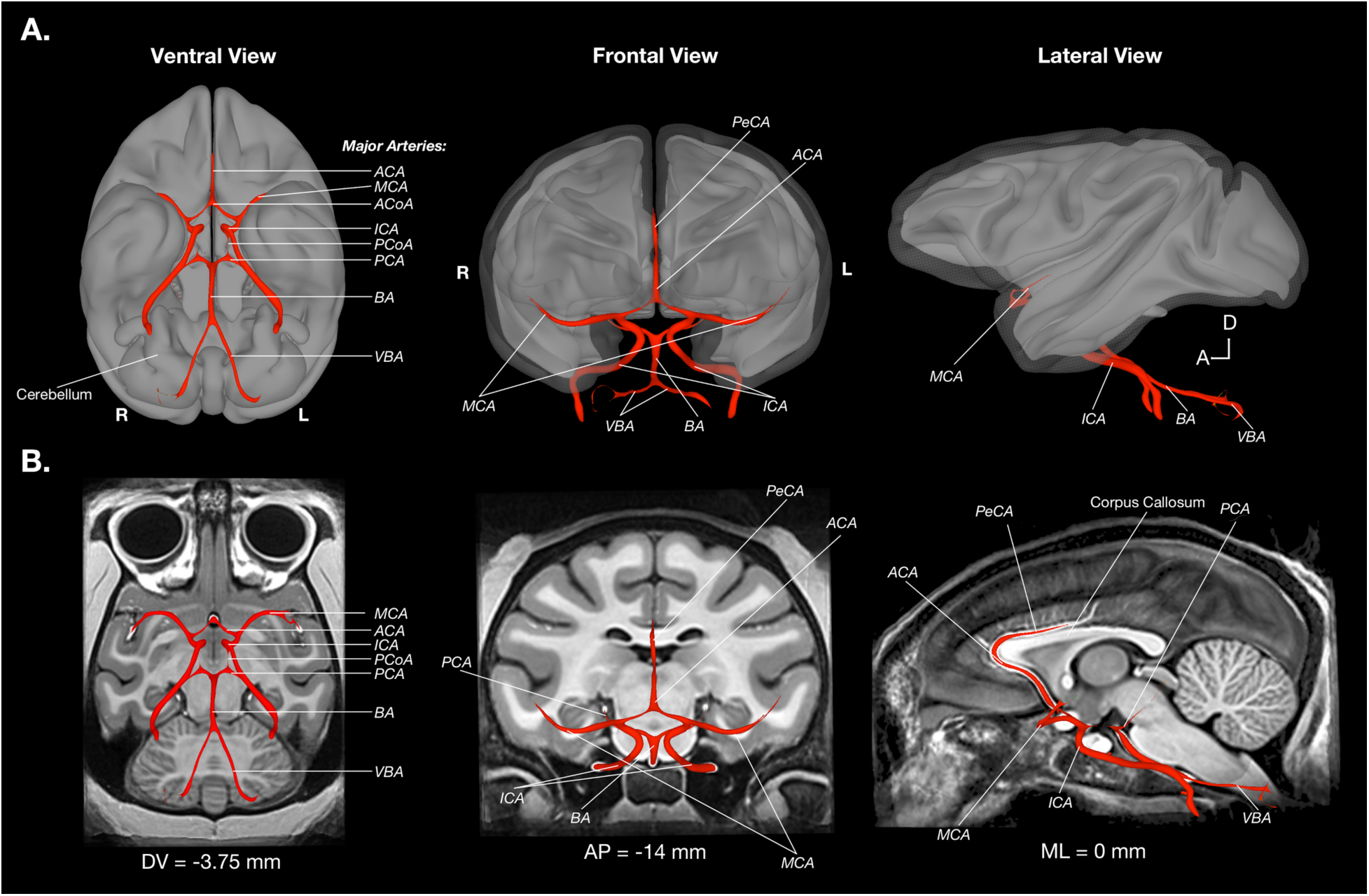
Surface visualization of the NMT arterial blood vasculature. To depict the topography of the arterial blood vasculature (BV), the arteries are shown as a surface overlay on a background of the 3-dimensional NMT GM and WM surfaces (A) as well as example 2-dimensional NMT slices (B). BV was approximated by selecting voxels from the raw image with intensity values above an arbitrary threshold, followed by manual editing and surface modeling. By consulting known atlases of macaque vasculature, all major brain arteries were identified in the NMT volume. The BV mask and surfaces are included in the NMT distribution. *ACA* = Anterior cerebral artery, *ACoA* = anterior communicating artery, *BA* = basilar artery, *ICA* = internal carotid artery, *MCS* = middle cerebral artery, *PCA* = posterior cerebral artery, *PCoA* = posterior communicating artery, *PeCA* = pericallosal artery, and *VBA* = vertebral artery.

### 3.8 Morphological Variability

Figure 6 shows the Mean Positional Difference (MPD) map on the “mid-cortical surface” of the NMT. The MPD represents the average displacement (in mm) of a given voxel in the NMT to the respective voxel location in each subject. Overall, the MPD values were quite low, with a brain-wide average of 0.45 mm (*i.e.*, ~1 voxel at the resolution of the original scans), and a maximum of 1.07 mm. There was no significant effect of gender, weight, or age as covariates on MPD in a voxel-wise 1-sample t-test (3dttest++ command) after correction for multiple comparisons (FDR, *q* < 0.05). The lack of a significant effect may be due to the affine alignment prior to the calculation of the nonlinear diffeomorphic transformations to the NMT; this prior transformation would largely nullify the effect of gender dimorphisms in brain size (Scott et al., 2015).

The NMT cohort’s MPD results are similar to estimates of morphological variability reported in previous macaque and baboon templates (Black et al., 2004; McLaren et al., 2009; Frey et al., 2011; Love et al., 2016). MPD was greatest in ventrolateral prefrontal cortex (VLPFC), anterior temporal cortex, and portions of occipital, and medial parieto-occipital cortex. This is consistent with previous work showing that areas of the VLPFC (as well as orbitofrontal cortex), anterior temporal lobe, and the occipital lobe have high variability in structure across individuals (Carmichael & Price, 1994; Van Essen et al., 1984; Maunsell & Van Essen, 1987).

### 3.9 NMT Cortical Thickness

In order to assess the structural characteristics of the NMT, we generated a map of cortical thickness (CT), shown in Figure S2. The pattern of CT was similar across both hemispheres. CT tended to decrease along the rostral-caudal dimension, with thicker estimates (~3-4 mm) found in subregions of frontal and rostral temporal cortex. Gyral portions of the occipital cortex, especially medially, were also relatively thick. Thinner estimates (~1-2 mm) were observed deep within sulci (e.g., caudal parts of the lateral and cingulate sulci, and central sulcus). These results are quite similar to the pattern of CT estimates found using a T2-weighted *ex vivo* template (Calabrese et al., 2015), as well as in earlier work (Koo et al., 2012). These results help validate the NMT as a representative anatomical volume, with typical macaque cortical morphometry.

### 3.10 Automated Single-subject Analysis

Processing the anatomical scan of a single monkey to generate a brain mask, classify tissue types, and characterize cortical morphometry is currently a largely manual process that is time consuming and requires knowledge of both brain anatomy and MRI analysis tools. The NMT eliminates the need to repeat this process for each new monkey as the anatomy can be nonlinearly transformed to the NMT volume, where these masks (as well as surfaces) have already been constructed and characterized. There is also the option to display data from individual subjects on the high quality and representative NMT anatomy.

However, some users may prefer to perform analyses in the subject’s native space and display results on the original anatomy. For example, this approach might be used by those who wish to avoid the interpolation of functional or anatomical voxels involved in the registration process. To facilitate analysis in a subject’s native space, we developed a novel, automated single-subject structural MRI processing pipeline. This pipeline leverages the tools available for the NMT dataset to generate masks for new subjects, thus eliminating the time-consuming step of hand drawing masks of different tissue types for each slice of an anatomical volume.

Table S1 describes our processing scripts for single-subject structural MRI analysis. *NMT_subject_align* employs the nonlinear registration method from previous work (Reveley et al., 2016), *NMT_subject_process* is designed for template-based brain extraction and tissue segmentation, and *NMT_subject_morph* estimates cortical morphometry. These scripts call on AFNI and ANTs commands to align the single subject’s brain to the NMT, then use the NMT’s masks as priors in generating masks for the subject, and then transforms these masks back to the subject’s native space.

Starting with a raw reconstructed T1-weighted anatomical volume (including the skull), the single-subject volume is first registered to the NMT using a combination of linear and nonlinear transformation procedures (Reveley et al., 2016). Prior to segmentation, the single-subject volume is corrected for non-uniform image intensities using the *N4BiasFieldCorrection* command (Avants et al., 2011). Brain extraction is then performed with the *antsBrainExtraction.sh* command, using the brain mask from the NMT as a prior (Avants et al., 2010). The *antsAtroposN4.sh* script, which iterates between the *N4BiasFieldCorrection* and *Atropos* commands, then segments the brain mask into three probabilistic tissue categories - CSF, GM, and WM - using the NMT probabilistic segmentation maps as priors (Avants et al., 2011).

Cortical thickness is estimated as in section 2.7 using the NMT’s cortical GM mask, nonlinearly warped to the individual subject. An option to estimate other morphometric features (e.g., surface area and curvature) is included as well. These features are estimated using the *SurfaceCurvature* command, a volume-based method which estimates curvature within a local neighborhood around a given voxel (Avants & Gee, 2003).

### 3.11 Accessing the NMT Dataset and File Structure

The NMT anatomical volume is publicly available and can be downloaded from the following links: <u>https://afni.nimh.nih.gov/pub/dist/atlases/macaque/nmt</u> and <u>http://github.com/jms290/NMT</u>. Also available for download are the brain mask (Figure 3a), the probabilistic tissue segmentation masks for GM, WM, and CSF, 4-tissue segmentation (inclusive of BV within the brain mask), and separate masks of the cortical GM, BV, olfactory bulb, and cerebellum. These volumetric data are stored in the NIfTI-1 (<u>https://nifti.nimh.nih.gov/nifti-1</u>) file format within a 253 × 347 × 245 voxel grid. Surfaces are distributed in GIFTI format (<u>http://www.nitrc.org/projects/gifti/</u>), with separate left and right hemisphere files. Additionally, the three scripts necessary for single-subject analysis (*NMT_subject_align* for registration, *NMT_subject_process* for generating the brain mask and tissue segmentation, and *NMT_subject_morph* for morphometric analysis) and accompanying documentation are provided. Finally, transformations to and from other templates are provided. Instructions for use of these files and citing this work can be found on the websites listed above.

## Discussion

We have produced a new multi-subject MRI template of the macaque brain (the NMT), coupled with visualization resources and automated analysis tools. The NMT contains anatomical detail superior to previous *in-vivo* monkey templates (Black et al., 2004; McLaren et al., 2009; Quallo et al., 2010; Van Essen et al., 2011; Frey et al., 2011; Rohlfing et al., 2012) and comparable to that of *ex vivo* templates (Calabrese et al., 2015; Reveley et al., 2016). These *ex vivo* templates do not reflect the natural and complete morphology (e.g, the olfactory bulb) of experimental subjects scanned *in vivo* because the tissue is fixed, lacking blood, and not encased in dura and skull. Moreover, these *ex vivo* templates also used different contrasts than the typical *in vivo* T1-weighted anatomical scan collected on most subjects.

The quality of the NMT volume makes it possible to resolve subcortical structures (see Figure 4), cortical laminae in some areas, fine substrates such as dura mater, and many arterial blood vessels. The NMT’s detailed anatomy derives in part from the ANTs nonlinear template generation procedure (Avants et al., 2010), the number of contributing subjects, and the higher acquisition field strength of our scans relative to other templates. In addition, the up-sampling of the resolution of the original scans (from 0.5 mm to 0.25 mm isotropic voxels) permitted finer-scale alignment of the 31 independent subjects. Because of these factors, the NMT contains emergent properties that were not visible or less pronounced in individual subject scans. We anticipate that still finer detail and alignment could be achieved by including additional subjects during template creation and acquiring their anatomical scans at higher resolution.

### 4.1 Applications of the NMT

The NMT volume, and accompanying segmentation masks and surfaces, can be used in a variety of applications. For example, the NMT volume and surfaces make striking backgrounds against which to present any kind of neuroimaging data (e.g., functional, structural, or diffusion imaging). One might also present whole-brain multi-subject functional MRI activations on the NMT’s mid-cortical and inflated surfaces. Activations could be further described in terms of their spatial coordinates in NMT space, which have the benefit of being quantitative, anatomically precise, and not subject to the vicissitudes of nomenclature. Activations can also be described in terms of the brain areas involved, either by referring to a paper atlas or by nonlinearly warping one’s preferred digital atlas to the NMT using the tools we provide. The NMT can also be used with anatomical scans to, for example, characterize the extent of a lesion, as well as with non-imaging data to, for example, indicate the target of tracer injections or electrode penetrations.

For most purposes, the NMT’s fine-tuned brain mask and combined segmentation map can be applied to data nonlinearly warped to the NMT. We provide a script (*NMT_subject_align*, outlined in Table S1) to register individual T1-weighted anatomies to and from the NMT brain, thus eliminating the need for the time-consuming steps of skull-stripping and segmenting tissue classes for individual subjects. Using the transformations from *NMT_subject_align*, the NMT’s CSF or BV mask could be used to control for non-neuronal sources of resting state correlations; the NMT’s cortical GM mask could also be used to confine analyses to voxels that are contiguous on the cortical sheet. These capabilities should streamline analysis for researchers conducting non-human primate imaging research.

Certain analyses, however, are specific to the individual’s anatomy or require precision in the subject’s native space. In these cases, it is preferable to tailor the brain mask and tissue segmentation to the individual subject. We provide a script (see *NMT_subject_process* in Table S1) that uses the NMT’s maps as a starting point to improve the accuracy of single-subject segmentation. This approach could be especially relevant for looking at differences (e.g. cortical volume) between groups of individuals, or for performing analyses (e.g., masking) at the single subject level. This tool could also be particularly useful for surgical planning or invasive procedures that target specific areas, such as the placement of individual electrodes or cortical arrays, as well as the injection of histological tracers, pharmaceutical agents, or optogenetic vectors. For investigations of brain structure for individual subjects, we also provide a script (see *NMT_subject_morph* in Table S1) for estimating different features of brain morphometry (i.e., CT, surface area, and curvature). This might be useful for comparisons with functional or behavioral data, where the dimension of interest is at the level of the individual subject rather than at the level of the group.

The NMT package is of benefit to those performing analysis of imaging data at either the single-subject or group level. We have linked the NMT to additional resources. For example, we provide nonlinear transformation matrices to the F99 template and associated anatomical atlases within the Caret software package (Van Essen et al., 2011). We also provide transformations to the D99-S volume and anatomical atlas (Reveley at al., 2016). The NMT thus establishes a foundation upon which to easily combine and analyze data from many subjects and sources. Bringing different kinds of data (e.g., functional, structural, and diffusion imaging) to the NMT’s standard coordinate space will facilitate comparison across imaging modalities as well as with data obtained through other methodologies (e.g., stimulation, inactivation, electrophysiology, EEG).

Furthermore, the template simplifies coordination and collaboration between researchers and groups with expertise in these different techniques. For example, several labs have identified regions of the macaque ventral temporal stream involved in processing faces, body parts, objects, curvature and color (Bell et al., 2009; Pinsk et al., 2009; Ku et al., 2011; Lafer-Sousa et al., 2013; Tsao et al., 2003, 2008; Moeller & Freiwald 2008; Yue et al., 2014). The NMT not only provides a rigorous basis for assessing the consistency of these visual category maps across subjects, but it also provides a means to assess the underlying morphology of areas selective for different characteristics. Conducting research with the NMT’s web of resources and tools should not only make it easier to understand one’s own data but also simplifies cross-subject, multi-modal, and collaborative research.

## Conclusion

We present the NMT, an anatomical template of the macaque brain, derived from 31 monkeys. The NMT also includes tissue maps, surfaces, and transformation scripts to assist in data analysis (<u>https://afni.nimh.nih.gov/pub/dist/atlases/macaque/nmt</u>). As non-human primate imaging progresses, the topics of scientific inquiry grow more ambitious. Answering such challenging questions will likely require more subjects and verification, which may entail combining data across research groups. We believe that the NMT can properly support collaborations to answer important questions, such as the variance of cortical topographies across macaques, the consistency of functional connectivity across a population, the longitudinal trajectories of brain morphometry, and the structural and functional homologies between human and non-human primates. By providing an open and universal platform for data visualization, characterization, and analysis, the NMT will assist researchers pursuing these challenging lines of inquiry, and further our understanding of the primate brain.

## Acknowledgments

The authors would like to thank David Yu for his help in the scanning procedures and data collection, as well as Alex Clark for his help in collating the demographic information and the raw image files. We also thank researchers in the Laboratory of Brain and Cognition and the Laboratory of Neuropsychology who contributed anatomical scans of their subjects to this project. This work was funded by the Intramural Research Program of the NIMH (ZICMH00289). The authors report no conflicts of interest.

